# *SleepMat*: A New Behavioral Analysis Software Program for Sleep and Circadian Rhythms

**DOI:** 10.1101/2022.01.31.478545

**Authors:** Shiju Sisobhan, Clark Rosensweig, Bridget C Lear, Ravi Allada

## Abstract

*Drosophila* is a premier model for the study of circadian rhythms and sleep, revealing molecular mechanisms that are conserved with vertebrates, including humans. The Drosophila Activity Monitoring (DAM) system has been employed for automated, high throughput analyses, enabling measures of a wide range of measures of circadian and sleep, including period, rhythmicity, anticipation, sleep amount, bout length, and bout number. Yet there are no publicly available programs that capture the full breadth of these measures. Here we develop a new program *SleepMat* that analyzes a uniquely broader range of circadian and sleep parameters. The program is built on a Matlab platform, employs an easy-to-use graphic user interface, and is highly flexible to customize data inputs. This program will enable a user-friendly high throughput analysis of a broad range of sleep and circadian parameters that can be coupled to the power of *Drosophila* genetics.

## 1. Introduction

Circadian clocks govern nearly all 24 *hr* rhythms in behavioral, physiological, cellular, and molecular processes in all organs and tissues (Allada & Chung, 2010; Patke, Young, & Axelrod, 2020). Sleep is a major output of the circadian clock and though sleep is a highly conserved process in both vertebrates and invertebrates, its function is poorly understood. The fruit fly, *Drosophila melanogaster*, has been a valuable model organism for the study of both circadian rhythms and sleep. Its utility stems in part from its simple genetics and the vast array of tools developed by the fly community. One such tool, the Drosophila Activity Monitor (DAM) system (TriKinetics, Waltham, MA), uses infrared beams to automate tracking of locomotor behavior rhythms. Flies are placed individually into a glass tube that is bisected with an infrared beam. The DAM system records the number of infrared beam breaks in each tube as a measure of locomotor behavior. A single DAM monitor has 32 channels and data is collected in 1 min bins and stored as a text (.txt) file. Thus, these small devices allow for cheap, easy, and massively parallel tracking of fly locomotor behavior with broad applications for the sleep and circadian fields.

From the simple monitor data collected by the DAM, a large variety of metrics for sleep and circadian studies can be calculated. Circadian analytics are relatively straightforward. Circadian rhythms can be quantified by their amplitude or strength of rhythmicity, periodicity, and phase under constant conditions. Activity counts can be binned and graphed as eductions or they can be shown as actograms. Periodicity can be determined from chi-squared periodogram analysis. More complex analysis is devoted to anticipation indices, which assess when and how well the flies anticipate lights on and lights off under light-dark conditions. Sleep, in contrast, requires more in-depth analysis. Sleep in *Drosophila* consists of a sustained period of immobility and meets several criteria set out for sleep: it is regulated by a homeostatic mechanism, behavioral immobility is accompanied by an increased arousal threshold, and the condition is reversible by strong stimuli (Campbell & Tobler, 1984; Hendricks et al., 2000; Shaw, Cirelli, Greenspan, & Tononi, 2000). From a data analytics perspective, sleep in *Drosophila* is defined as uninterrupted behavioral inactivity lasting for 5 min. This definition is based on initial experiments describing sleep in flies (Shaw et al., 2000). Flies exhibiting normal locomotor activity readily respond to a vibration stimulus of moderate intensity. However, flies that have been behaviorally inactive for 5 min or longer show a reduced behavioral response to these stimuli (Hendricks et al., 2000; Shaw et al., 2000; van Alphen & van Swinderen, 2013). Thus, initial analysis of DAM data requires scanning for >5 min epochs of immobility. From this initial analysis, sleep can be further characterized by its duration as well as its architecture including sleep latency (how quickly one falls asleep), initiation (sleep bout number), and maintenance (sleep bout length). Moreover, many sleep experiments include a perturbation step (sleep deprivation) and the amount of sleep lost during this perturbation and gained following it (sleep rebound) can be calculated. Mathematical calculation and biological interpretation of these sleep parameters are provided in the result section. Comparison and further study of these parameters is critical for sleep and circadian research; hence it is very important to calculate sleep and circadian parameters easily, accurately, efficiently, and cost-effectively. Additionally, due to the large throughput of *Drosophila* experiments, scalable, comprehensive analysis through a simple interface is critical.

Current software packages do not integrate analyses of the full spectrum of circadian and sleep parameters. Existing software for analyzing DAM system data are ShinyR-DAM(Cichewicz & Hirsh, 2018), PySolo (Gilestro & Cirelli, 2009), SCAMP (Donelson et al., 2012), Counting Macro (Pfeiffenberger, Lear, Keegan, & Allada, 2010), Rethomics (Geissmann, Garcia Rodriguez, Beckwith, & Gilestro, 2019), and RhythmicAlly (Abhilash & Sheeba, 2019). These programs provide only a limited number of sleep and circadian parameters; thus, the user must rely on multiple software suites with unique input files to get all parameters, which is time-consuming and error prone. Table-I summarizes the various sleep parameters which are commonly used in sleep analysis, and it shows which parameters are calculated by the existing software. Moreover, some software does not provide a user interface, so the user must perform analysis from the command line every time. Here we introduce ‘*SleepMat*’ (sleep analysis software based on MATLAB) software implemented entirely in MATLAB with a user-friendly graphical user interface (GUI) to analyze *Drosophila* activity monitoring data. It is straightforward to use and can calculate more than 25 sleep and circadian parameters within a short time, which will reduce the user time and effort considerably.

**Table I.** Comparison of *SleepMat* with existing software.

| Variables | ShinyR-dam<br>(Cichewicz & Hirsh, 2018) | PySolo<br>(Gilestro & Cirelli, 2009) | SCAMP<br>(Donelson et al., 2012) | RhythmicAlly<br>(Abhilash & Sheeba, 2019) | Counting macro<br>(Pfeiffenberger et al., 2010) | Rethomics<br>(Geissmann et al., 2019) | SleepMat |
| --- | --- | --- | --- | --- | --- | --- | --- |
| Sleep | Yes | Yes | Yes | Yes | Yes | Yes | Yes |
| Bout number & bout length | Yes | Yes | No | No | Yes | Yes | Yes |
| Total activity | Yes | No | Yes | Yes | Yes | Yes | Yes |
| Activity/waking min | Yes | Yes | Yes | Yes | Yes | No | Yes |
| Latency | No | Yes | No | No | No | Yes | Yes |
| Morning & evening anticipation | No | No | No | No | No | No | Yes |
| Morning & evening anticipation phase | No | No | No | No | No | No | Yes |
| Sleep loss & gain | No | Yes | Yes | No | No | No | Yes |
| Latency after sleep deprivation | No | No | No | No | No | No | Yes |
| Eduction results | Yes | No | No | No | Yes | No | Yes |
| Actogram | Yes | Yes | Yes | Yes | No | Yes | Yes |
| Periodogram | Yes | No | No | Yes | No | Yes | Yes |
| Number of Triage conditions | 1 | 0 | 1 | 0 | 2 | 1 | 3 |
| Lifespan | No | No | No | No | No | Yes | Yes |
| Triage channel | Yes | No | Yes | No | Yes | Yes | Yes |

## 2. The software

*SleepMat* is developed in MATLAB version 2019a on a Windows platform and modified for the Mac operating system. Both Windows and Mac versions also function well with the 2021b version. We have created two type of SleepMat software which helps those who have MATLAB installed and those who don’t. A) Standalone application, which doesn’t require Matalab installed on the system B) p-code that can run SleepMat in MATLAB, which requires a MATLAB license. SleepMat software and the instructions to install the standalone application are available on the GitHub repository (https://github.com/shijusisobhan/SleepMat2022.1). Download the appropriate folder based on the operating system. Steps to run *SleepMat* in MATLAB are described as follows:

1. Open MATLAB
2. Change the current folder to the new folder where *SleepMat* software is located. E.g., If *SleepMat* software is saved in the ‘D’ drive, then to change the current folder, type the following line in the MATLAB command window:

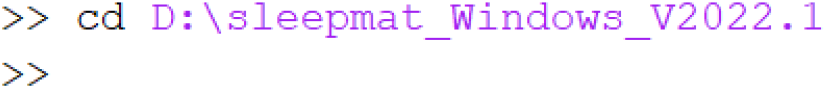
3. To open the software, type ‘sleepmat’ in the MATLAB command window, then hit enter.

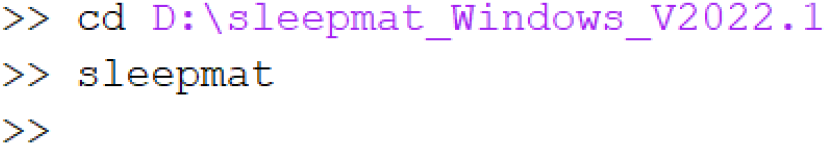
4. A window (GUI) will open as shown in Fig 1.

**Fig 1.**
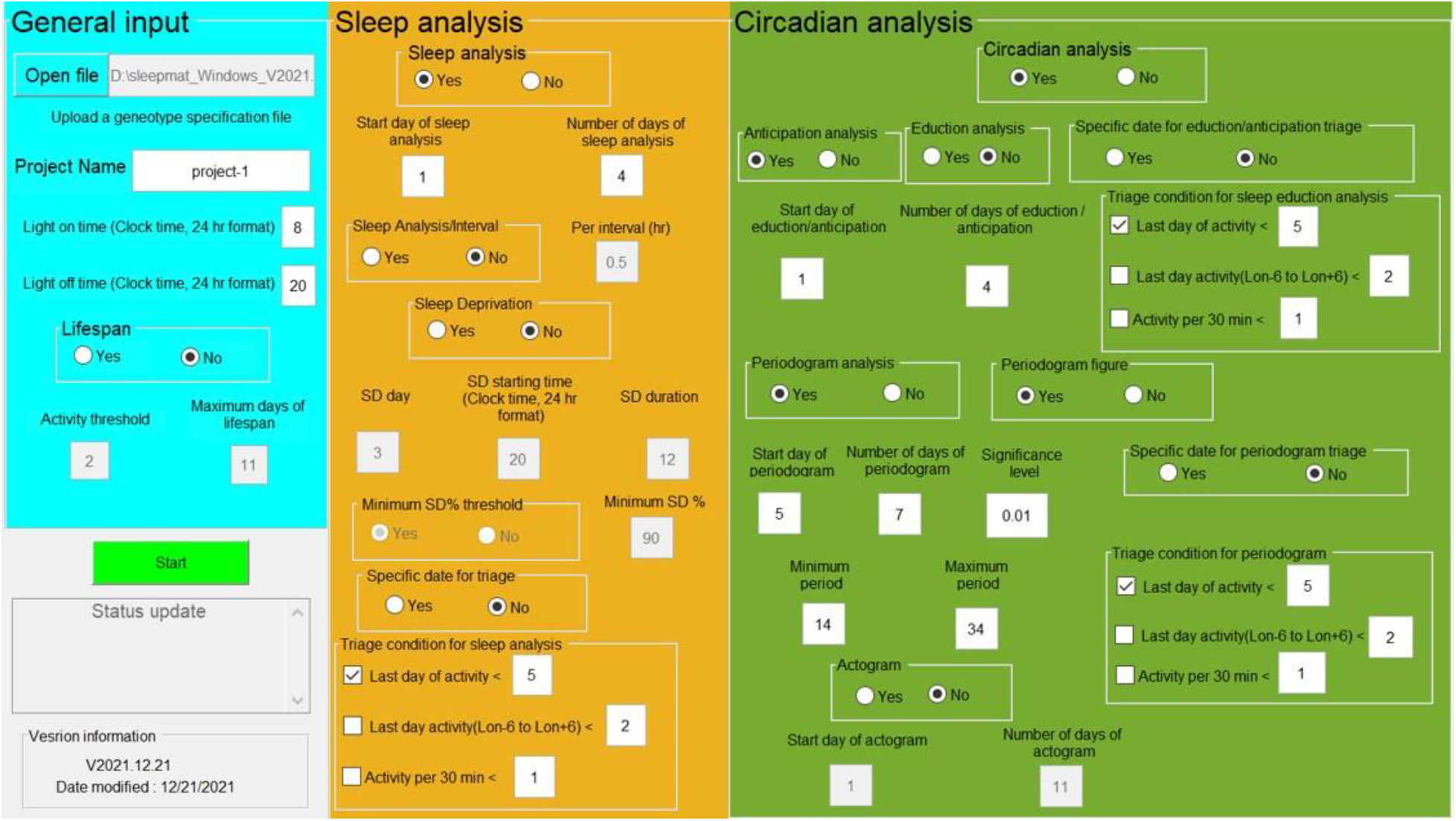
*SleepMat* graphical user interface: The graphical user interface (GUI) contains three panels. The general input panel is used to upload genotype information, enter the project name, input daily lights-on and lights-off times, activity threshold and maximum lifespan days. The sleep analysis panel contains inputs for sleep and sleep deprivation analysis. The circadian analysis panel contains inputs for anticipation and periodogram analysis, as well as production of eduction, periodogram, and actogram figures. Users can select/deselect each analysis. Triage conditions for different analyses are also included in each subsection.

### 2.1 Monitor files and genotype specification file

S*leepMat* accepts DAM system monitor files with data in one min bins. The files should not be processed by DAMFileScan. A genotype specification file, comprised of a simple Excel file, describes the details of the experimental conditions and links to the appropriate monitor file. Steps to arrange monitor files and format genotype specification files are described here:

1. Place all the monitor files recorded by the TriKinetics DAMSystem3 data acquisition software into a single folder (Fig 2). The file name should be ‘Monitor#.txt’, where # is a numeric value that indicates the board number. Note: If processing multiple runs with an overlapping set of monitor numbers, rename one of the monitor files by, for instance, appending the run number along with the monitor number. **E.g.,** Board number 32 was used in runs 1845 and 2970, so both runs require a file named Monitor32.txt by default. The monitor file from run 1845 can be renamed as Monitor184532.txt and the associated monitor number can be changed in the genotype specification file described below. This approach allows an unlimited number of runs to be processed at once.
2. Create an excel spreadsheet (.xlsx) for the genotype specification file in the same folder as the monitor files and give it any file name. A sample format for the genotype specification file is given in Fig 2.
3. Format your genotype specification file:

3.1. First Row A1-F1 (Mandatory): These are the header fields. You can give them any name, as long as there is text in the first row. Note that column G is optional.
3.2. Column A (Mandatory): The run number should be entered here and can be numeric or alphanumeric. The first cell (A1) is the header and from A2 onwards are the run numbers. Any arbitrary alphanumeric value can be recorded here to differentiate each independent experiment, but the cell cannot be left blank.
3.3. Column B (Mandatory): The monitor number should be entered here and must be a numeric value. E.g., For the file ‘Monitor102.txt’, 102 should be entered in this column.
3.4. Column C (Mandatory): The genotype name should be entered here, and it can be either alphanumeric or plain text. Note: the forward slash character (/) cannot be used in the genotype name.
3.5. Column D, E (Mandatory): These columns encode the range of channels for analysis. Column D is the start channel and column E is the end channel for any given genotype. These are numeric values and should be between 1 and 32, corresponding to loading positions in Trikinetics’ *Drosophila* Activity Monitors. Note: if a genotype runs across multiple boards, each board’s flies must be entered in a separate line. However, all flies from the same genotype will be grouped in the final analysis.
3.6. Column F (Mandatory): The start date of analysis is entered here and its format is the same as what appears in the monitor file (e.g., 13 Jan 18). It must be plain text.
3.7. Column G (Optional): A specific triage date can be entered here as an optional setting. To evaluate triage conditions on a specific date, enter the date here. Date format is the same as what appears in the monitor file (e.g., 16 Jan 18). If this column is left blank, by default the software will evaluate triage conditions using the last day of analysis.

**Fig 2.**
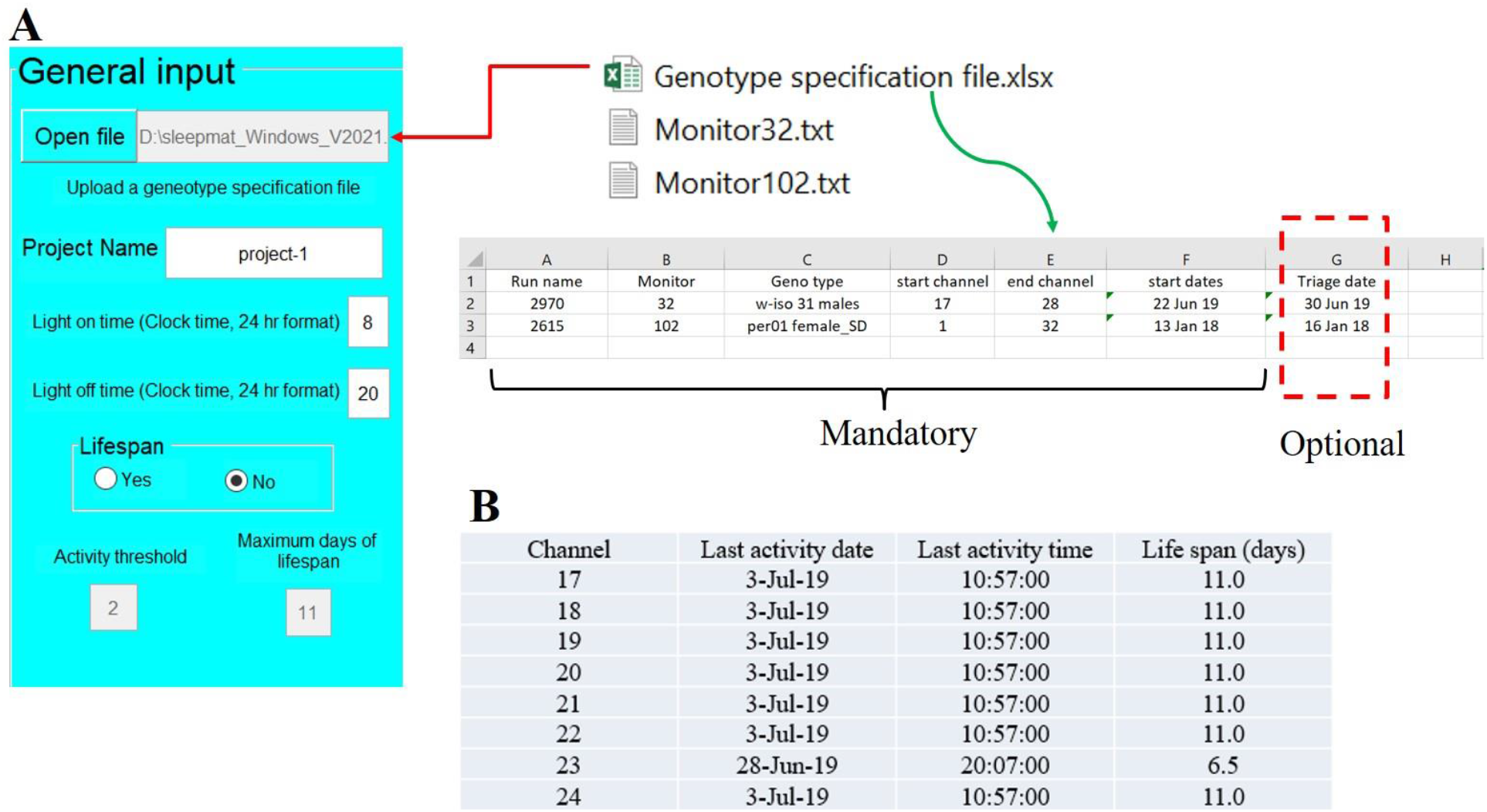
General input and genotype specification file: A) Top: user clicks ‘Open file’ button to browse and upload genotype specification file. In the genotype specification file, the user specifies the genotype name, monitor files, channels, analysis start date, and alternative triage date (optional) for the analysis. Project name: user enters a project name for the analysis. All results will be stored in a folder with this project name. Light offset and onset: user enters lights-on and lights-off times (based on DAM clock time) to indicate the beginning of the day and night in light:dark (LD) conditions. Bottom: user enters activity threshold and maximum days for lifespan. B) Lifespan results. *SleepMat* records the date and time of the last activity of each fly and calculates lifespan in days.

### 2.2. General Input

The first panel of the *SleepMat* GUI is the general input where the user can upload the genotype specification file and enter light on/off times, and lifespan information. Detailed descriptions for general input items follow:

1. **Open file**: Click the ‘Open file’ button to browse and select the genotype specification file for the analysis.
2. **Project Name**: Users can assign a project name for the current analysis. The program automatically creates a folder with the project name in the current working directory and saves all the results there.
3. **Light on time (Clock time, 24hr format)**: This is the light onset time of the experiment. The default value is 8 AM. Note that it is specified in clock time with 24 *hr* format. **E.g.,** If the light onset time is 8 AM, then enter 8, if it is 1 PM, then enter 13.
4. **Light off time (Clock time, 24hr format)**: This is the light offset time of the experiment. The default value is 8 PM, which is specified in clock time as 20.
5. **Lifespan (Yes/No)**: Increasingly *Drosophila* is used as a model to link sleep and circadian rhythms to longevity (Hendricks & Sehgal, 2004). *SleepMat* calculates the number of days that a fly is alive based on an activity threshold. Users can select to perform lifespan analysis by checking Yes.
6. **Activity threshold**: This is the minimum value of activity to determine the lifespan of a fly. To find lifespan, total activity over 24 *hr* is computed in a 1-min moving window manner (e.g. from 22-Jun-19 8:00:00 to 23-Jun-19 8:00:00, from 22-Jun-19 8:01:00 to 23-Jun-19 8:01:00 and so on). If the total activity over a window is less than the activity threshold, then life span is calculated based on the starting time of that window. E.g., if the activity from 28-Jun-19 20:06:00 to 29-Jun-19 20:06:00 is less than the threshold, then the last activity is recorded as 28-Jun-19 20:06:00, and the life span is calculated in days (Fig 2B).
7. **Maximum days of lifespan**: If DAM boards are left plugged into the system beyond the scope of the experiment, the DAM system will continue to collect more data. The user may not be interested in the additional data and can exclude those days from lifespan analysis by setting a maximum threshold. If the lifespan exceeds the maximum days entered here, *SleepMat* automatically assign lifespan as ‘Maximum days of lifespan’.

## 3. Results

### 3.1. Sleep Analysis

Sleep in *Drosophila* is defined as uninterrupted behavioral inactivity (0 activity counts) lasting for 5 min or more. Therefore, any bout which has greater than or equal to five consecutive zeros (with 1 min activity bins) is considered a sleep bout. Total sleep in an interval is the total number of zeros in all sleep bouts within that interval (Fig 3). Bout length in minutes is the number of consecutive 1 min bin zeros in a sleep bout. Average bout length is the total sleep divided by total bout number (see Fig 3). Longer bout length is often an indication of deeper sleep (van Alphen & van Swinderen, 2013); on the other hand, shorter bout length and high bout number is an indication of fragmented sleep. Sleep latency is the number of minutes until the beginning of the first sleep bout after lights off (Fig 3). Calculating sleep latency is the best way to quantify difficulty initiating sleep (Eiman & Harbison, 2019). Total activity is the sum of all non-zero values in an interval. Waking activity (activity/waking min) is the total activity divided by the total number of non-zero values (Fig 3). It is an indicator of physical activity levels and has been used to distinguish sick flies (reduced waking activity) from long sleepers (normal waking activity). It can also be used to distinguish short sleepers with normal waking activity from hyperactive short sleepers.

**Fig 3.**
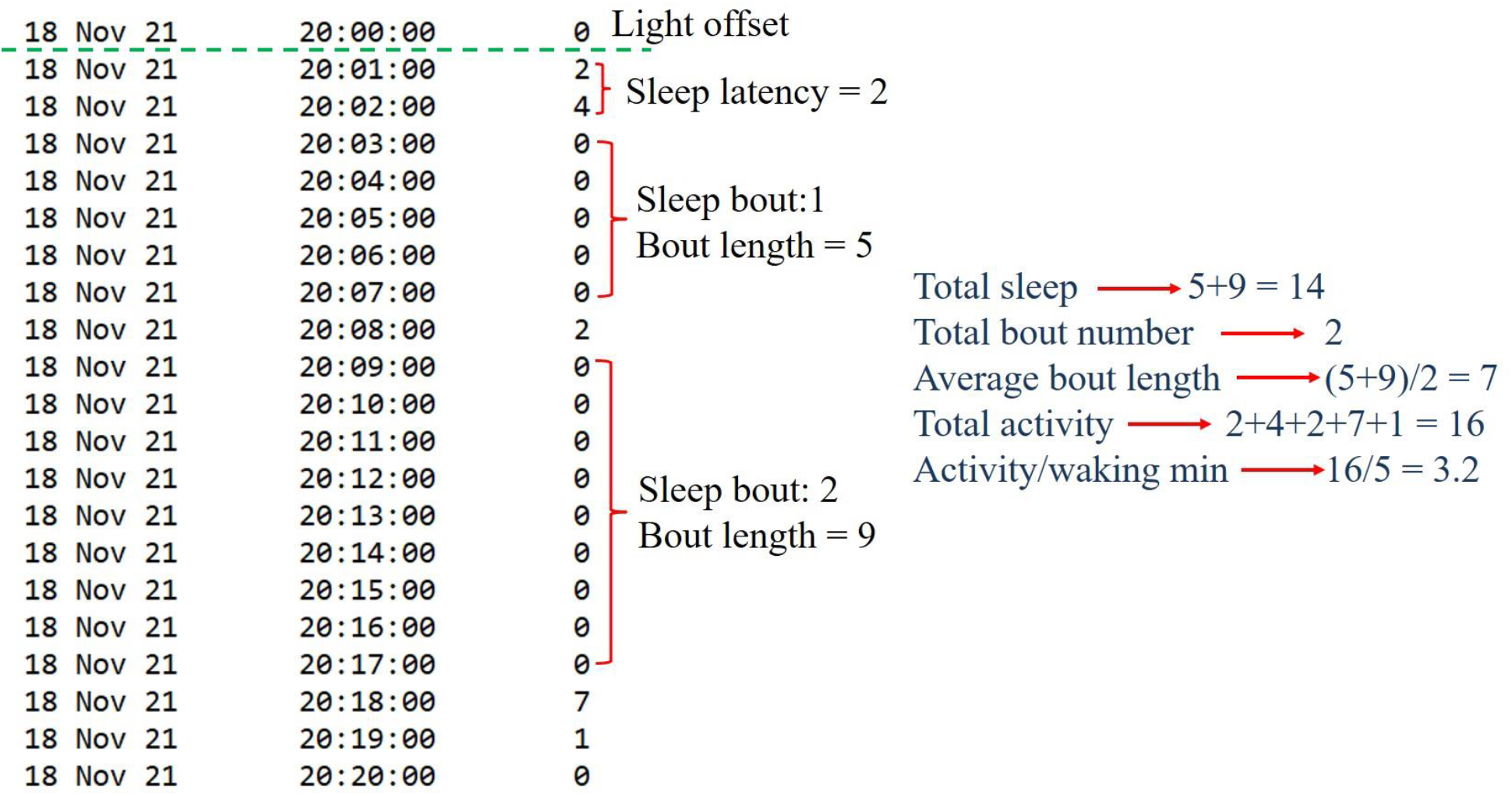
Sleep parameter and activity calculations. Each number represents a record of the number of times a single fly breaks an infrared beam in one minute. The green dashed line represents the lights off time. Sleep is defined as complete inactivity for a duration of 5 minute or more. Non-zero numbers represent activity.

Steps to calculate aforementioned sleep parameters are described as follows:

1. **Sleep analysis (Yes/No)**: Users can select to perform sleep analysis by checking Yes.
2. **Start day of sleep analysis**: Enter the starting day of sleep analysis. The default value is 1. If the start day of sleep analysis = 1, then the sleep parameters will be determined starting from the ‘start date of analysis’ specified in the genotype specification file. The start day of sleep analysis cannot exceed the number of days of data present in the data file. Start time for the analysis corresponds to the lights-on time.
3. **The number of days of sleep analysis**: Enter the number of days to analyze. The default value is 4. **E.g.,** if the start day is 13 Jan 18 and the number of days of analysis is 4, then analysis starts on 13 Jan 18 at 8:00 (light on time) and ends on 17 Jan 18 at 8:00 (Fig 4B). The number of days of analysis should not extend past the end of the data available in the data file.
4. **Sleep analysis/Interval (No/ Yes)**: Users can specify an interval by first selecting Yes, then entering the desired interval for analysis. Sleep parameters evaluate on this specified interval. The default is No.
5. **Per interval (hr)**: If the user changes the sleep analysis/interval option to Yes, then the user can enter a desired interval. The default is 0.5 (30 min). **Eg.** For 30 minutes, it would break down every 30 minutes and average over the total number of days. (8-8:30 AM, 8:30-9 AM, etc).

**Fig 4.**
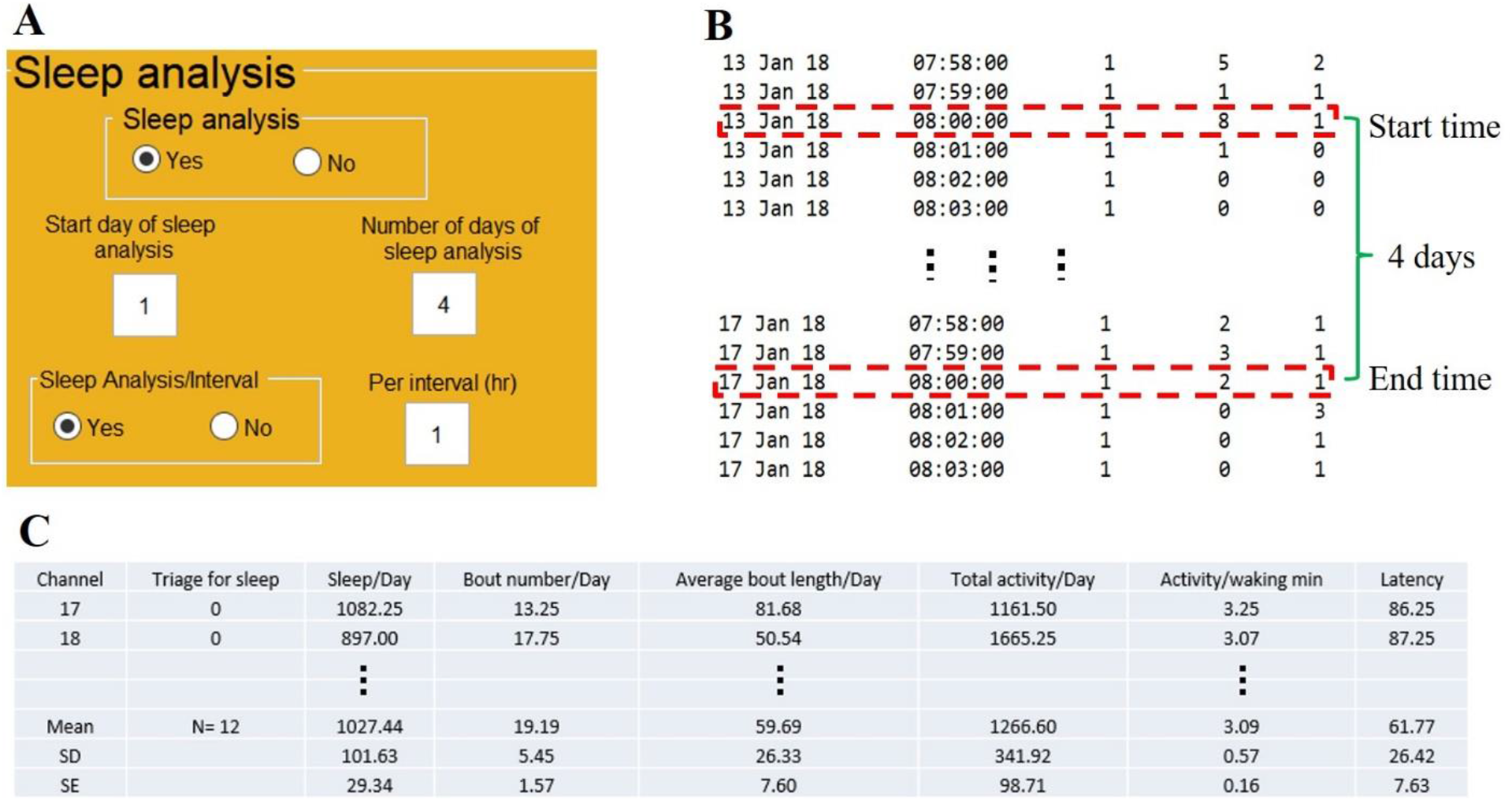
Sleep analysis: A) Input panel for sleep analysis. User enters the start day, the number of days of analysis, and the interval for sleep analysis. B) Example of sleep analysis range as determined from DAM file information and user inputs. If the start day of sleep analysis entered is 1, then the analysis begins from the start date of analysis specified in the genotype specification file (‘13 Jan 18’ in this example). Start time for the analysis is based on the lights-on time specified in the general input panel (first red dashed box; 8:00 in this example). One day of analysis is the 24 *hr* period between lights on times. The second red dashed box indicates the end time of 4 days of sleep analysis, as specified in the sleep analysis panel (A). C) Sleep analysis results. The program reports total minutes of sleep per day, the number of sleep bouts per day, average bout length, total activity per day, activity per waking min, and sleep latency for each fly as well as the mean, standard deviation (SD) and standard error of the mean (SE) for each genotype.

Fig 4. describes the input panel for sleep analysis (Fig 4A), data selection for sleep calculations (Fig 4B), and the sleep and activity parameters that are calculated by *SleepMat* (Fig 4C). By default, *SleepMat* calculates sleep per day (24 *hr*). However, the user can select an interval for sleep calculations in which *SleepMat* calculates sleep over that interval. For each individual, the sleep and activity parameters are calculated and stored in excel files. *SleepMat* also reports averages over all individuals in a genotype and calculates the standard deviation (SD) and standard error of the mean (SE) for each genotype.

#### Sleep deprivation analysis

Sleep deprivation in flies is typically followed by an increase in the amount and duration of sleep (sleep rebound). To quantify sleep loss and sleep gain (rebound), a specific deprivation or rebound window is compared with a baseline window. These measures are important to study the effects of sleep deprivation to better understand homeostatic regulation of sleep. A typical example of a sleep deprivation experiment is shown in Fig 5B. Calculation of sleep loss and sleep gain is given as:

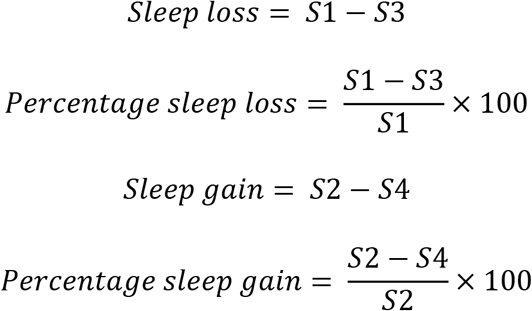

**Fig 5.**
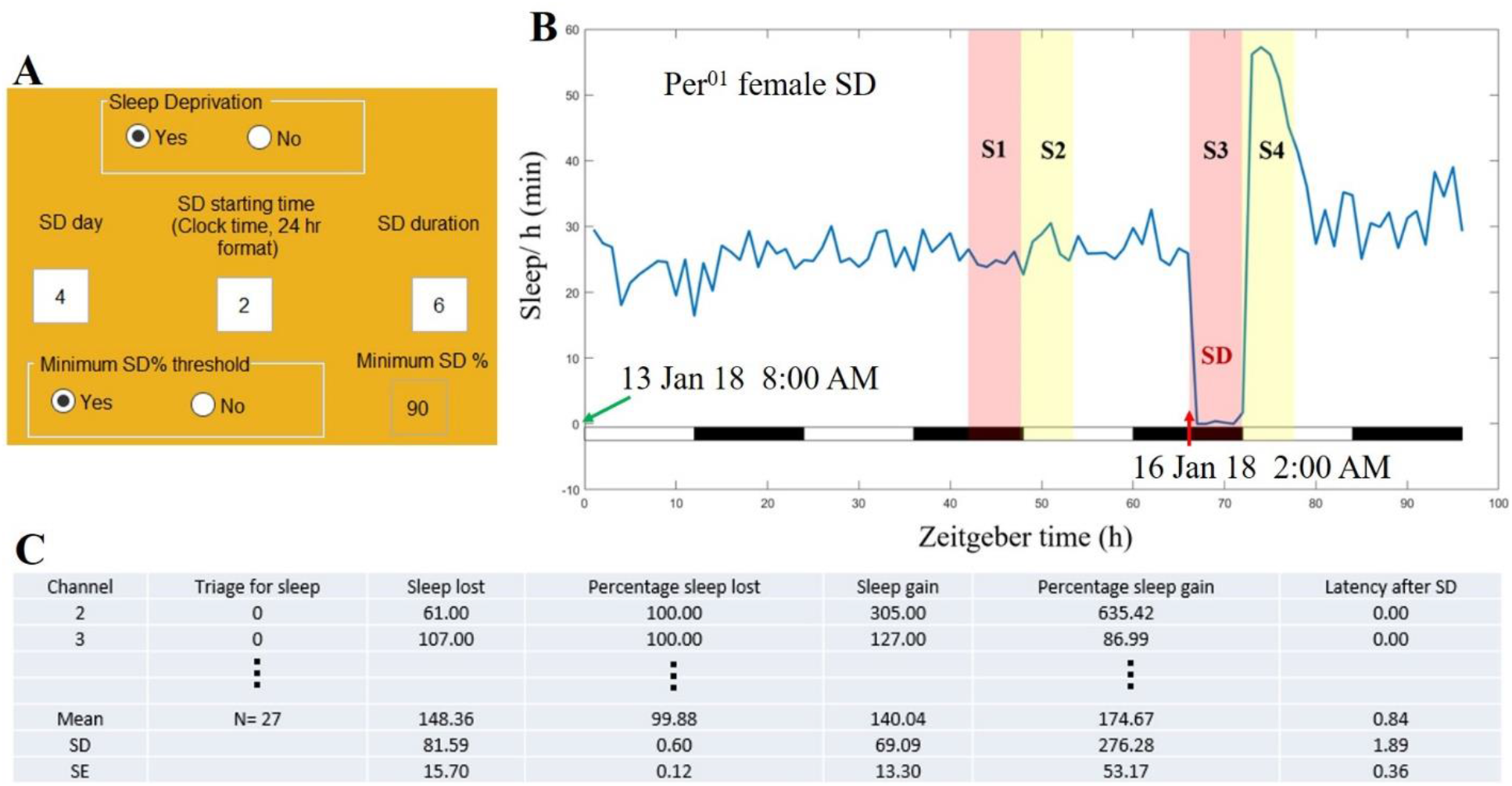
Sleep deprivation analysis: A) Input panel for sleep deprivation analysis. User enters the day, start time, and duration of sleep deprivation (SD). Start time is based on the DAM file clock, not Zeitgeber time. The ‘Minimum SD% threshold’ option sets a threshold value to determine whether the flies are sufficiently sleep deprived. If the sleep loss exceeds this threshold, then only those flies are considered sleep deprived. B) Illustration of 6 *hr* sleep deprivation. The black bars at the bottom indicate the dark phase and the white bars indicate the light phase. The green arrow indicates the starting time of analysis and the red arrow indicates the starting time of sleep deprivation. In this example, day, start time, and duration of sleep deprivation are 4, 2, and 6 respectively (16 Jan 18 from 2:00 AM to 8:00 AM). Baseline sleep (S1) is measured from 2 AM to 8 AM on the day prior to deprivation (15 Jan 18), followed by a 6 *hr* measurement after this baseline period (S2; 8 AM to 2 PM on 15 Jan 18). Total sleep is also measured during the sleep deprivation period (S3), and over the 6 *hr* just after the sleep deprivation (S4; 8 AM to 2 PM on 16 Jan 18). C) Sleep deprivation results. The program reports minutes of sleep lost during deprivation (S3-S1), minutes of sleep gained after deprivation (S4-S2), percentage of sleep lost/gained, and latency after sleep deprivation.

Where S1, S2, S3 and S4 are described in the Fig 5.

Some researchers are also interested in sleep latency after sleep deprivation which is the number of minutes until the first sleep bout after sleep deprivation. *SleepMat* calculates sleep loss/gain, percentage of sleep loss/gain, and latency after sleep deprivation. Fig 5 shows the input panel for sleep deprivation analysis (Fig 5A), the sleep loss and sleep gain calculation (Fig 5B), and sleep deprivation analysis results (Fig 5C). Settings to perform sleep deprivation analysis are described as follows:

1. **Sleep deprivation (Yes/No)**: Sleep deprivation analysis can be performed by selecting Yes. The default is No.
2. **SD day**: The day when the flies are sleep deprived is entered here. The default is 3. This day is specified with respect to the start day of analysis. **E.g.,** if the start day of analysis is on 13 Jan 18, and sleep deprivation is on 15 Jan 18, then SD day is 3.
3. **SD starting time (Clock time, 24 hr format)**: This is the start time of sleep deprivation. The default is 8 PM (20).
4. **SD duration**: This is the duration of sleep deprivation. The default is 12 *hr*.
5. **Minimum SD% threshold (Yes/No)**: When analyzing sleep deprivation data, it may be desirable to only include flies that were deprived of a certain percentage of their sleep in the final analysis. This option is used to specify a threshold for inclusion. Only those flies whose sleep loss exceeds this threshold are considered sleep deprived and included in the final analysis. The default is Yes.
6. **Minimum SD%**: The minimum threshold value to count for sleep deprivation is entered here. It is a percentage value, and the default is 90.

### 3.2. Circadian analysis

Research on circadian rhythms often requires estimating period, power, activity pattern, and activity peaks (anticipation/phase) under light-dark or other environmental cycles. *SleepMat* is equipped to analyze time-series data to estimate the period and its statistical power (chi-square periodogram (Sokolove & Bushell, 1978)), activity patterns (actogram, eduction plot), and entrainment parameters (anticipation index and anticipation phase (Harrisingh, Wu, Lnenicka, & Nitabach, 2007)).

Anticipation parameters in circadian rhythms are the measure of any activity change in anticipation of lights on or lights off. Anticipation measures are helpful to study how the circadian clock regulates activity amount and timing in advance of environmental transitions (e.g., light-dark) under entraining conditions (e.g., light) (Seluzicki et al., 2014). *Drosophila* shows activity peaks under entrainment conditions with one peak anticipating lights on (morning anticipation) and the other anticipating lights off (evening anticipation). An eduction plot is a plot of normalized average activity typically in 30 min bins over N days (Fig 6B) to visualize activity peaks in anticipation of light transitions. It compiles data from multiple days into a single day (Pfeiffenberger et al., 2010), and the mathematical formula of normalized average activity is given as follows:

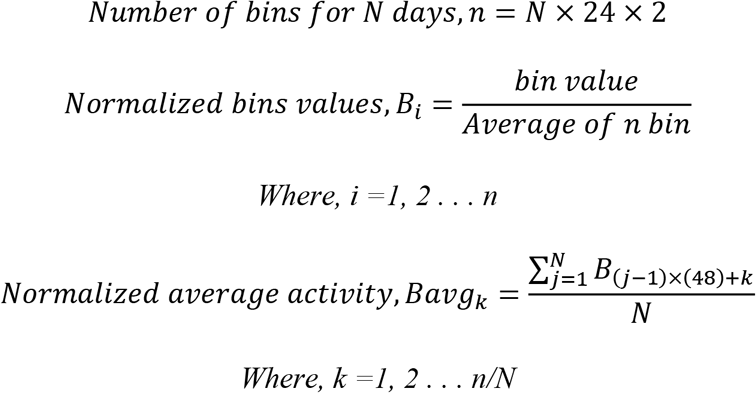

**Fig 6.**
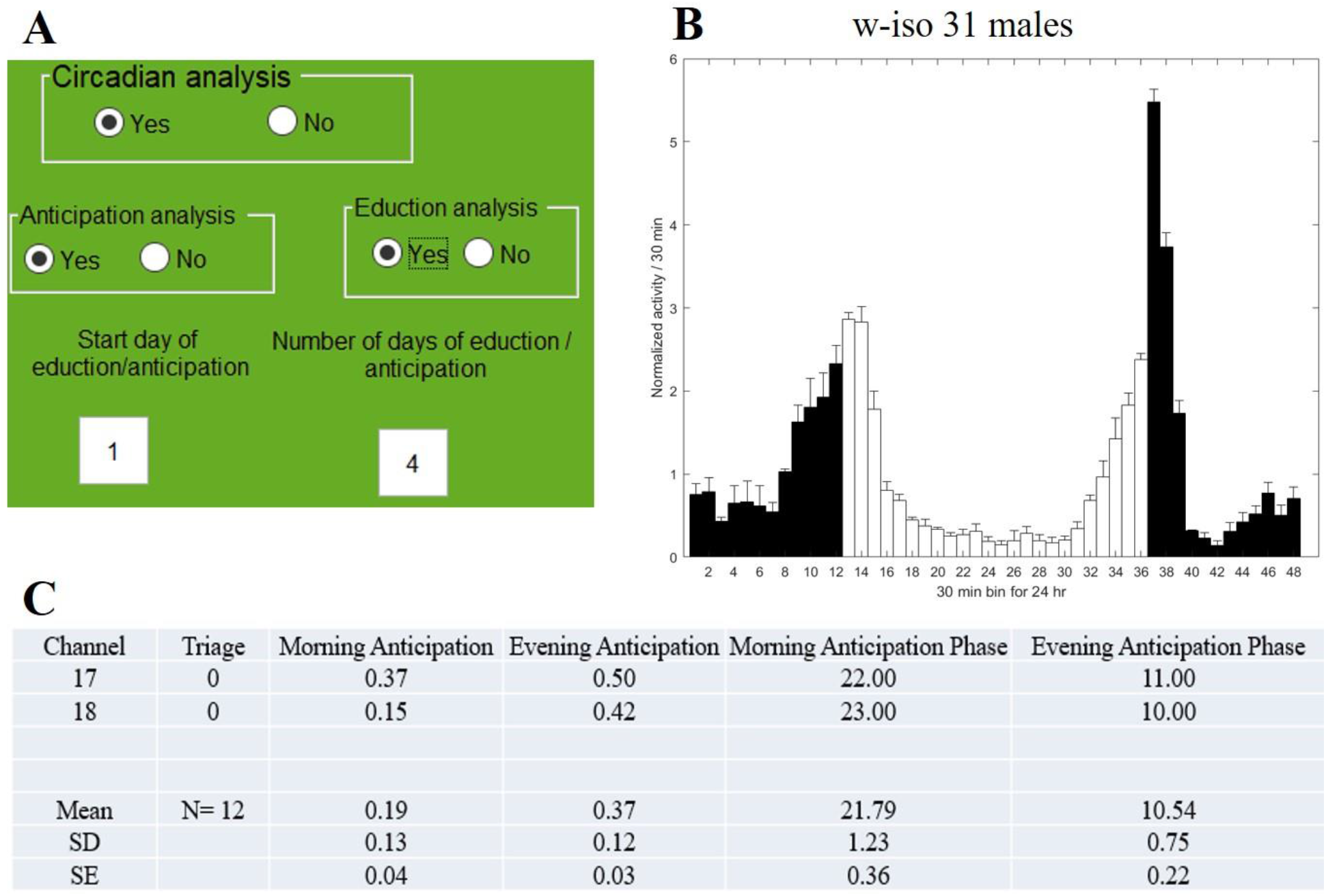
Eduction plot and anticipation analysis: A) Input panel for normalized activity plots (eduction) and anticipation analysis. Here we used a wild type iso 31 (w-iso31) male flies. Eduction plots and anticipation analyses are evaluated over the same days. The user enters start day and number of days of anticipation/eduction analysis. B) Example eduction plot. Black and white bins are 30 min normalized activity bins during dark and light phases respectively. Error bars represent standard error of the mean. Activity is averaged over all individuals in a genotype and over four days based on user input (A). C) Anticipation results. The program reports the amplitude and phase of morning and evening anticipation for individuals, and mean, standard deviation (SD) and standard error of the mean (SE) for each genotype.

Eduction plot assumes a 24 *hr* period and an offset of −6 *hr* and hence data for calculating eduction plots starts from 6 *hr* before the light on time. *SleepMat* estimates anticipation based on the method developed by (Harrisingh et al., 2007), in which the ratio of activity during the three hours before lights on/off and the activity during the six hours before lights on/off is presented as the anticipation index. The mathematical formulae for calculating anticipation measures are given as follows:

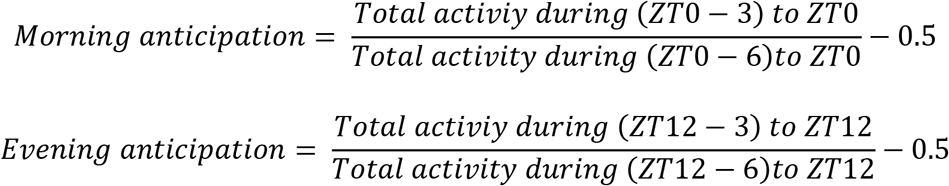

Anticipation phase:

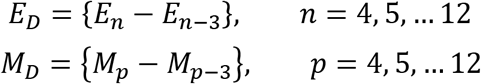

Where *E_n_* is the *n^th^* 30 min activity bin after *ZT12-6*, and *M_p_* is the *p^th^* 30 min activity bin after *ZT0-6*.
*Evening anticipation phase* = *ZT6.5* + (*n_max_*/2 – 1). Here *^n_max_^* is the value of *n* where *E_D_* is Maximum
*Morning anticipation phase = ZT18.5* + (*^p_max_^*/2 – 1). Here *p_max_* is the value of *p* where *M_D_* is Maximum

Fig 6 describes the input panel for eduction/anticipation analysis (Fig 6A), sample eduction plot and (Fig 6B), and anticipation results. Steps to estimate the eduction plot and anticipation parameters are given as follows.

1. **Anticipation analysis (Yes/No)**: Select Yes to perform anticipation analysis.
2. **Eduction analysis (Yes/No)**: Used to produce normalized activity profiles (eductions) for each genotype. Note: anticipation calculations and eduction analyses (if selected) will use the same analysis range.
3. **Start day of eduction/anticipation**: Enter the starting day of anticipation/eduction analysis. The default value is 1.
4. **Number of days of eduction/anticipation analysis**: Enter the number of days to be analyzed for eduction/anticipation.

#### Periodogram

*SleepMat* computes the period of *Drosophila* circadian rhythms and determines the statistical significance of the computed period based on the chi-square periodogram procedure (Enright, 1965; Sokolove & Bushell, 1978). Time resolution for the chi-squared periodogram is set to 30 min and the significance line is calculated with an adjustable alpha (significance level) value. Fig 7 describes the input panel for periodogram estimation (Fig 7A), sample periodogram visualization (Fig 7B), and periodogram results. Steps to estimate the periodograms are described as follows:

1. **Periodogram (Yes/No)**: Select Yes to perform chi square periodogram analysis for individual flies.
2. **Periodogram figure (Yes/No)**: If the user would like to produce an average periodogram figure for each genotype, then select Yes.
3. **Start day of periodogram**: Enter the start day for periodogram analysis.
4. **Number of days of periodogram**: Enter the number of days of periodogram analysis.
5. **Significance level**: Here the user can set the desired alpha level for detection of periodicity using chi-squared periodiogram analysis.

**Fig 7.**
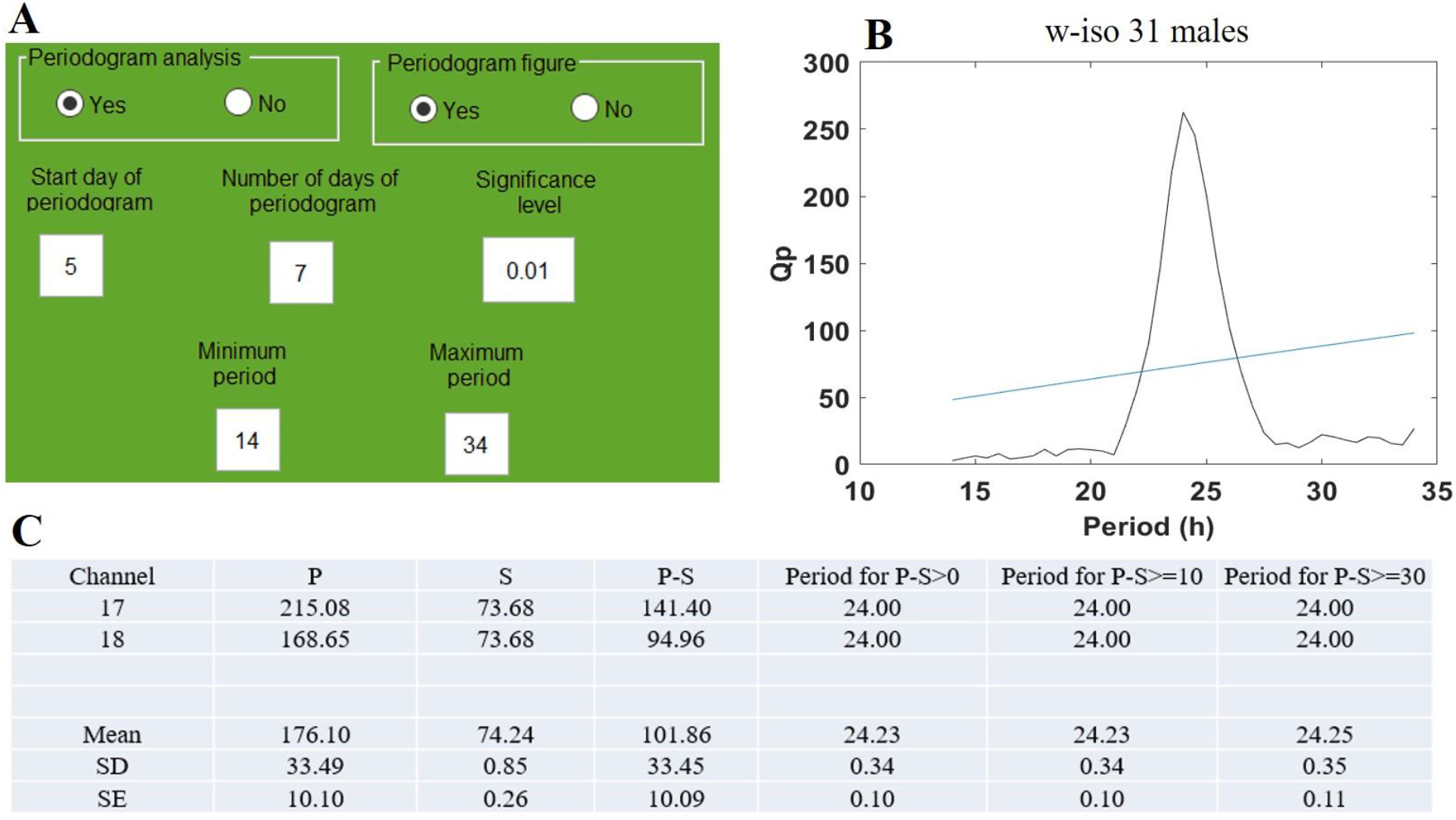
Periodogram: A) Input panel for periodogram analysis. Analyses are performed using chi-squared periodogram, typically calculated from constant conditions. The user enters the start day and number of days for periodogram analysis. The minimum and maximum periods for consideration and threshold for significance can be adjusted as needed. B) Example periodogram figure. The period being tested plotted along the x-axis and power along the y-axis. The blue line is the significance line calculated based on the significance level given in the input panel. Power is averaged over all individuals in a genotype. The time resolution of the periodogram is 30 min. C) Individual periodogram results. For individuals with a Power value (P) exceeding the significance value (S), Power, Significance, and P-S difference are reported. Period values are also reported using three different thresholds: all individuals with P-S>0, P-S >10, and P-S >30.

#### Actogram

Circadian locomotor activity rhythms are normally represented as a graph called an actogram, where horizontal lines represent days and thick black bars plotted on each horizontal line represent the activity. *SleepMat* produces a double plotted actogram, where the *x*-axis represents two days (48 *hr*). It plots two days on each horizontal line and shows the second day on the far right of each line, as well as at the beginning of the second horizontal line, and so on (Fig 8B). Steps to produce the actogram are described below:

1. **Actogram (Yes/No)**: To produce double-plotted actogram figures for individual flies, select Yes.
2. **Start day of actogram**: Enter the start day for the actograms that will be generated. The default is 1.
3. **Number of days of actogram:** Enter the number of days that will be represented on the actogram.

**Fig 8.**
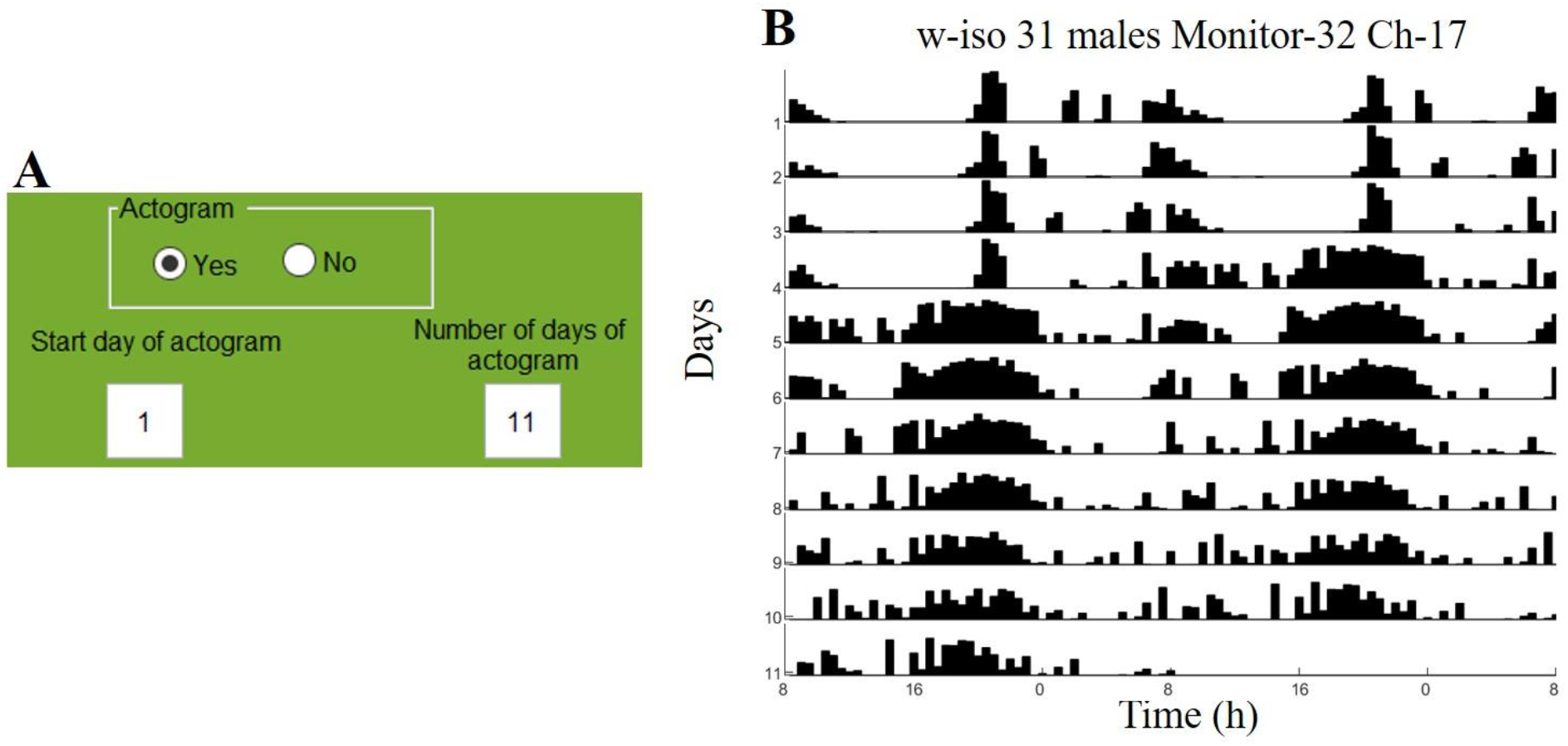
Actogram: A) Input panel for actogram. The user enters the start day and number of days in the actogram. B) Double plotted actogram. Two days are plotted on each horizontal line with the second day comprising the second half of each line, as well as the first half of the line below, and so on. Thick black bars plotted on the horizontal line represent the activity in each 30-minute bin.

### 3.3. Triage

One issue that analysis programs need to handle is the removal or triage of flies that die during the analytical period. Our software is flexible, accurately detecting dead flies based on any of the 3-triage conditions (Fig 9A). Triage results are given values of 1 for dead flies, and 0 for alive (Fig 9B). Dead flies are automatically removed from the analysis. Information about the number of active flies in each genotype is recorded in the excel file. Steps to determine the triaged flies are described as follows:

1. **Specific date for triage (Yes/No):** If the user wishes to evaluate triage conditions on a specific date, then select Yes. When No is selected, *SleepMat* will triage on the last day of analysis. When Yes is selected, then the desired triage date must be included in Column G of the genotype specification file.
2. **Triage condition for sleep/eduction/periodogram analysis**: This is the condition for determining whether the fly is dead or not. There are 3 triage conditions that are commonly used by different sleep research groups. An explanation for each triage condition is given below:

a. **‘Last day activity < ##’**: If on the last day (or the triage date specified in the genotype specification file) a fly exhibits fewer activity counts than threshold value, then the fly is treated as dead. We note that lights-on and -off can trigger the infrared emitter detector pairs and thus result in anomalous counts. Hence, we set the default threshold as 5.
b. **‘Last day activity (Lon-6 and Lon+6 < ##)’**: If the sum of the 6 *hr* activity before light onset and the 6 *hr* activity after light onset on the last day (or the day you specified on the genotype specification file) is less than threshold value, then that fly is considered dead. We set the default threshold as 2 to consider the anomalous count due to the light-on.
c. **‘Activity per 30 min < ##’**: This condition evaluates over the number of days of analysis. Thus, if a fly’s activity counts per 30 min are on average (over the evaluation period) less than the threshold, then that fly is treated as dead. Default threshold value set as 1, which can also be used to exclude extremely low activity flies (Pfeiffenberger et al., 2010). For example, a fly with an activity/30 min reading <1 had <192 total activity counts for 4 days of the behavior run.

**Fig 9.**
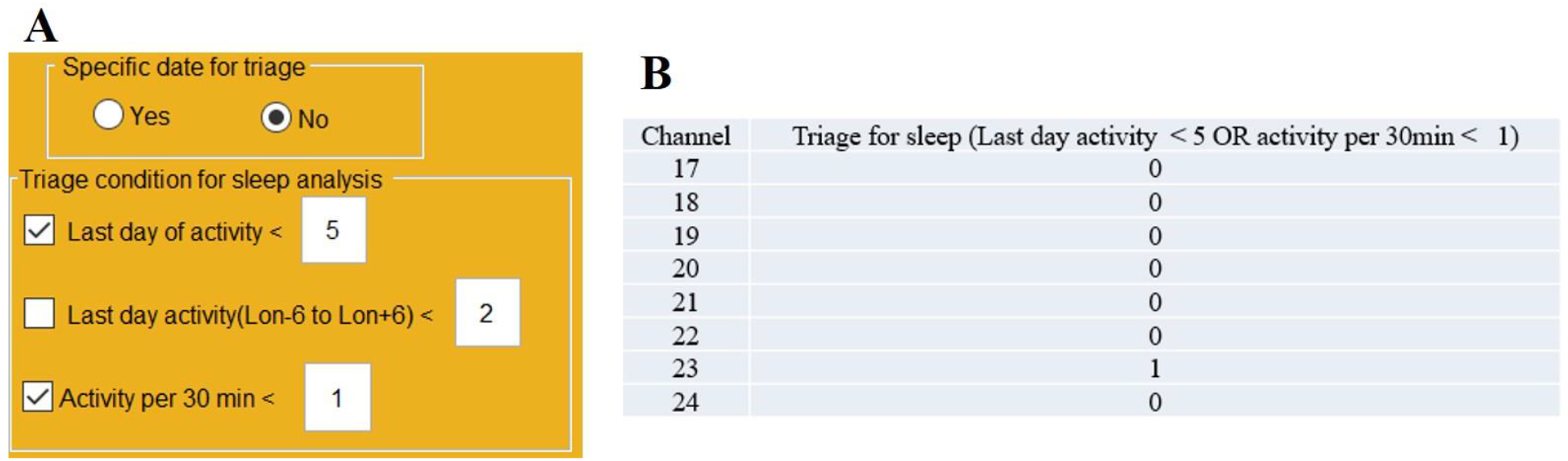
Triage: A) Input panel for triage condition. If the user wishes to evaluate triage conditions on a specific date, select ‘Yes’ and enter that date in the genotype specification file. Default is No, and it will evaluate triage on the last day of analysis. Users can choose a triage condition and threshold values of activity for determining whether flies should be removed from analysis. B) Triage result. Accurate detection of dead flies based on any of the 3 triage conditions. Triage value 1 indicates a dead fly and 0 indicates alive.

## 4. Discussion

*Drosophila* is a preeminent model for understanding the molecular and genetic mechanisms of sleep and circadian rhythms. The DAM system records daily locomotor behavior from *Drosophila*. In this work, we introduce *SleepMat*, a MATLAB-based software, to analyze DAM system data. Compared with existing software, *SleepMat* is simple, fast, robust, and versatile with many novel features (Table I).

### Simplicity

The availability of the user-friendly GUI will attract researchers who are unfamiliar with coding. *SleepMat* does read “Monitor files” directly, therefore there is no need to use the DAMFileScan operation to cut the DAM files into “channel files”. The addition and removal of flies is easily accomplished by adding or removing a row in the genotype specification file. Also, there is no need to specify each fly individually; instead, each genotype is assigned to a range of channels. This straightforward input file helps to streamline analysis, especially when analyzing a large number of experiments with multiple genotypes and replicates.

### Fast and robust

*SleepMat* can calculate more than 25 sleep and circadian parameters within a short time with fewer mouse clicks. The results are automatically saved in Excel files and figures in .png format. In a single run, the user can analyze as much data as possible. It provides individual results and averages over all individuals, which enables the study of groups as well as interindividual variation. Triage conditions automatically detect the dead or poorly active flies and exclude them from the analysis, which considerably reduces the user’s time and effort and improves the validity of results.

### Versatile

*SleepMat* allows users to calculate sleep parameters for light (L) and dark (D) phases as they may be distinct processes. The periodogram feature allows adjustments of the range of tested periods and filters arrhythmic flies based on an adjustable significance level. Finally, the dead flies are identified based on three different triage conditions with an adjustable activity threshold, which provides flexibility to researchers to define dead or sick flies. *SleepMat* analysis is not limited to classic 12 *hr* photoperiod (12 *hr* day, 12 *hr* night), instead, the software can easily accept a different range of photoperiods by changing light on time and light off time. This appeals to researchers who are working with seasonal variations in circadian rhythms. *SleepMat* can perform separate data analyses for LD and DD days by setting start day and number of days analysis.

### Novelty

Many researchers are interested in quantifying the anticipation index (increases in activity before light transitions) and the anticipation phase (the time at which the largest increase occurs). Most researchers manually calculate anticipation and phase by using an Excel spreadsheet. One major novel feature of *SleepMat* is the automatic estimation of anticipation and phase which eliminates tedious manual calculation. Additionally, the eduction feature enables visualization of activity in the LD cycle. Besides identifying dead flies, *SleepMat* also provides information about the lifespan of each fly, ie. how long each individual is alive, which is a major novel addition compared to other software. Moreover, sleep deprivation analysis including latency after sleep deprivation is also a unique feature of *SleepMat*.

Given the power of *Drosophila* genetics and the high throughput nature of DAM activity analyses, *SleepMat* will provide an automated yet flexible way of analyzing DAM-based datasets across large scales, dramatically increasing the efficiency of such analyses. *SleepMat* will thus enable large scale analysis for sleep and circadian parameters in a single, simple workflow.

## Notes

https://github.com/shijusisobhan/SleepMat2022.1

### Competing Interest Statement

The authors have declared no competing interest.

